# Human innate lymphoid cell activation by adenoviruses is modified by host defence proteins and neutralizing antibodies

**DOI:** 10.1101/2022.06.13.495867

**Authors:** Océane Paris, Franck JD Mennechet, EJ Kremer

## Abstract

Innate lymphoid cells (ILCs), the complements of diverse CD4 T helper cells, help maintain tissue homeostasis by providing a link between innate and adaptive immune responses. While pioneering studies over the last decade have advanced our understanding how ILCs influence adaptive immune responses to pathogens, far less is known about whether the adaptive immune response feeds back into an ILC response. In this study, we isolated ILCs from blood of healthy donors, fine-tuned culture conditions, and then directly challenged them with human adenoviruses (HAdVs), with HAdVs and host defence proteins (HDPs) or neutralizing antibodies (NAbs), to mimic interactions in a host with pre-existing immunity. Additionally, we developed an *ex vivo* approach to identify how bystander ILCs respond to the uptake of HAdVs ± neutralizing antibodies by monocyte-derived dendritic cells. We show that ILCs take up HAdVs, which induces phenotypic maturation and cytokine secretion. Moreover, NAbs and HDPs complexes modified the cytokine profile generated by ILCs, consistent with a feedback loop for host antiviral responses and potential to impact adenovirus-based vaccine efficacy.

**Author Summary:** Several studies have shown the importance of innate lymphoid cells (ILCs) both from an immune and physiological point of view, in particular for their role in the maintenance of tissue integrity, pathogens clearance, or in the establishment of immune tolerance. Our study focuses on the role of ILCs during direct challenge with prototype vaccines based on human adenoviruses (HAdVs) ± host defence proteins (HDPs) or neutralizing antibodies (NAbs) to mimic interactions in a host with pre-existing immunity. In parallel, through an ex vivo approach we observe how bystander ILCs respond to the uptake of HAdVs ± NAbs by monocyte-derived dendritic cells. We show that ILCs take up HAdVs, which induces pro-inflammatory and antiviral responses through phenotypic maturation and cytokine secretion. Moreover, HAdV-NAb and HAdV-HDP complexes modified the cytokine profile generated by ILCs, consistent with a feedback loop for host antiviral responses and potential to impact HAdV vaccine efficacy.

## Introduction

Innate lymphoid cells (ILCs) are functional kin to CD4 T helper (Th) cells. In contrast to Th cells, ILCs traffic through the lymphatic and vascular systems to preferentially reside in mucosal compartments where they help maintain a balance between anti-pathogen immunity and tolerance [1–3]. Unlike T and B cells, ILCs do not express rearranged antigen-specific receptors [1]. ILC interactions with neighbouring cells are crucial events in the induction and development of immune responses [4, 5]. In synergy with myeloid cells, respond to pathogens through the secretion of cytokines [6, 7]. Like Th cells, ILCs can be functionally and phenotypically subdivided into subsets: ILC1 (which historically included cytotoxic NK cells), ILC2, and ILC3. NK cells appear to be counterparts of CD8^+^ T cells, while ILC1, ILC2, ILC3 the counterparts of Th1, Th2, Th17/22 CD4^+^ T cells, respectively [8, 9]. LTi (lymphoid tissue-inducer) cells belong the ILC3 family and are involved in embryonic lymph node formation. Yet, ILC subsets are not static and show context-specific heterogeneity and plasticity, particularly as we age and during the development of antiviral responses [10, 11].

By the time we are adolescents, we have been infected with several human adenovirus (HAdV) types [12, 13]. The archetypal robust and long-lived immune response against HAdVs is due, in part, to latent infections that persist for years and constantly re-stimulate the memory B and T cell responses [14–16]. HAdV are nonenveloped particles with a linear double-stranded DNA genome of ∼36 kilobase pairs. The more than 110 HAdV types are grouped into 7 species (A to G) [17]. The variable tropism of HAdVs typically causes mild, self-limiting symptoms within 10 days post-infection [18, 19]. Globally, HAdVs of species A and C mainly induce pathology in the respiratory, urinary and gastrointestinal tracts. Species B HAdVs infections have the broadest tissue diversity and can cause disease in the respiratory, urinary, gastrointestinal and conjunctiva [17, 18]. The species D HAdVs typically cause disease in the conjunctiva and gastrointestinal tracts, while those of species E affect the respiratory tract and conjunctiva. For HAdVs of species F and potentially G, symptoms are preferentially in the intestinal compartments [17].

In the era of COVID-19, HAdV-based vaccine efficacy and safety are of particular relevance [20–22]. The roles ILCs play against HAdVs and HAdV-based vaccines are unknown. Moreover, whether the responses by ILCs are affected by pre-existing HAdV immunity has not been addressed. To fill this gap, we evaluated the interactions between human ILCs and three HAdV types that are used as vaccines: HAdV species C type 5 (-C5), species D type 26 (-D26), and species B type 35 (-B35) [5,23,24]. These three HAdVs have different seroprevalence profiles and differ in the mechanism by which they are taken up by cells.

In this study, we initially tweaked a protocol for the culturing of ILCs from human blood. Then, we quantified ILC uptake of HAdV-C5, -D26, and -B35 alone, or in complex with host defence proteins (HDPs), or neutralizing antibodies (NAbs) [25–28]. We characterized the levels of potential HAdV receptors, receptors for HDP- and NAb-complexed HAdVs, and relevant pattern recognition receptors (PRRs). Finally, as ILCs cooperate with neighbouring antigen-presenting cells [6,29,30], we developed an *ex vivo* environment to mimic this interplay. We show that HAdVs complexed with HDPs or NAbs induced differential cell surface levels of activation markers, and production of pro-inflammatory and antiviral cytokines with activities comparable to that of Th cells [31]. As bystanders, the ILC response to monocyte-derived dendritic cells (moDCs) that are challenged with HAdVs ± NAbs, can be HAdV-type dependent. These data demonstrate that pre-existing B cell immunity against HAdVs and HDPs directly impact ILC responses, which likely influence vaccine efficacy.

## Results

### ILC purity and stability

ILCs were obtained by negative immunomagnetic selection from anonymous blood bank donor PBMCs. To evaluate ILC recovery and purity, we used multi-parameter flow cytometry and a combination of markers including Lin^-^, CD127^+^, CRTH2^+/-^ and CD117^+/-^. After enrichment, the cells were characterized according to their size and granulosity. Approximately 50% of the cells had a lymphoid profile (**Fig 1A)**. Within the lymphoid population, approximately 3% were CD3^+^ (**Fig 1B**), and approximately 60% were CD3^-^/CD127^+/-^ of which 22% were CD127^high^ (**Fig 1C**). In this donor, 28% of the cells were ILC1 (CRTH2^-^/CD117^-^), 16% were ILC2 (CRTH2^+^/CD117^+/-^), and 56% were ILC3 (CRTH2^-^CD117^+^) (**Fig 1D)**. Cumulative data from >60 donors highlight the heterogeneity of ILCs in anonymous blood bank donors (**Fig 1E)**. To identify non-ILCs in the enriched populations, we stained for NK, NKT, T, and B cells using CD16 and CD56, CD3, and CD19 and CD20, respectively. The percentage of contaminating NK and NKT cells was 0 - 5%, T cells 0 - 2%, and B cells 0.4 - 7% (**S1 Fig**).

**Fig 1.**
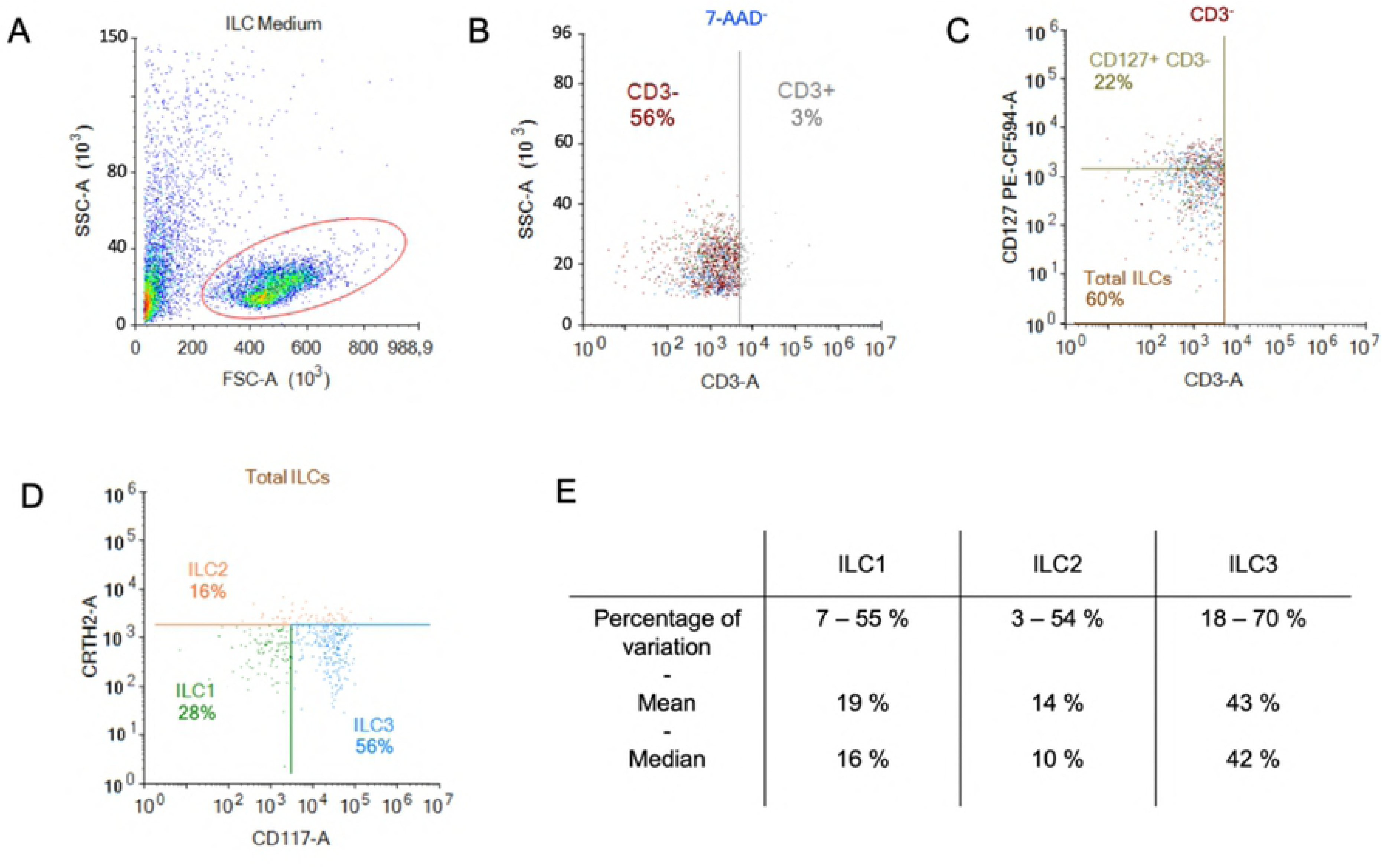
Enrichment and identification of ILCs from peripheral blood. Freshly isolated PBMCs were used for the negative selection of ILCs. **A)** Population of cells post-negative selection. Of these, ∼50% were lymphoid based on their size and granulosity;**B)** from the lymphoid population, we gated on the CD3^-^ population and **C)** in the CD3^-^ population the majority of cells were CD127^+^ (MFI for CD127^+^ was 2269 vs. 668 for CD127^-^)**; D)** from the CD3^-^ population we also screened for the presence of CRTH2 and CD117, which delimits the ILC populations; **E)** cumulative data from 60 donors showed the range of percentage, the median, and the mean of each ILC subset.

Initially, we maintained ILCs in RPMI/human AB serum/IL-7, but we were limited to assaying ILC phenotype and functionality during the first 24 h post-enrichment. To attempt to prolong this window, we tested “NK medium”, IL-2, and pyruvate. We found that the combination of RPMI/human AB serum (10%), IL-7 (10 ng/mL), and sodium pyruvate (1 mM) prolonged the phenotypic stability of the ILCs until ∼48 h post enrichment. Of note, IL-7 induces the internalisation of CD127, the α chain of the IL-7R. Therefore, from then on, we also gated on CD3^-^/CD127^low^ cells, followed by CRTH2 and CD117 to identify the ILCs.

### ILCs take up HAdV-C5, -D26 and -B35

Several cells are involved in the initial response to viral infections. ILCs could influence the immune response by responding to cells that take up viruses and/or by taking up the virus directly. To determine whether ILCs take up HAdVs, we incubated the cells with replication-defective (ΔE1) HAdV-C5, -D26, or -B35 vectors encoding GFP variants. At 24 h post-challenge, we found an average uptake efficacy of 13.5% for HAdV-C5, 13% for -D26, and 17% for -B35 (**Fig 2A**). We then broke these data down into the uptake of each HAdV type by each ILC subset. Globally, ILC2s take up all three HAdVs more efficiently than ILC1 & 3s (**Fig 2B-D**). The uptake of each HAdV for a given subset of ILCs, shows that ILC1 and ILC3s take up more HAdV-B35, followed by -C5 and -D26 (**S2 Fig**). ILC2s more readily take up HAdV-C5 and -B35. Each ILC subset thus shows a modestly variable uptake profile depending on the HAdV type. The notable difference in efficacy between donors (e.g., 0 - 80% of cells for HAdV-B35) is not unique to ILCs: primary cultures of monocytes and moDCs also show high interdonor variability [32–34]. Together, these data suggest that all ILC subsets could be involved in the detection of HAdV capsids.

**Fig 2.**
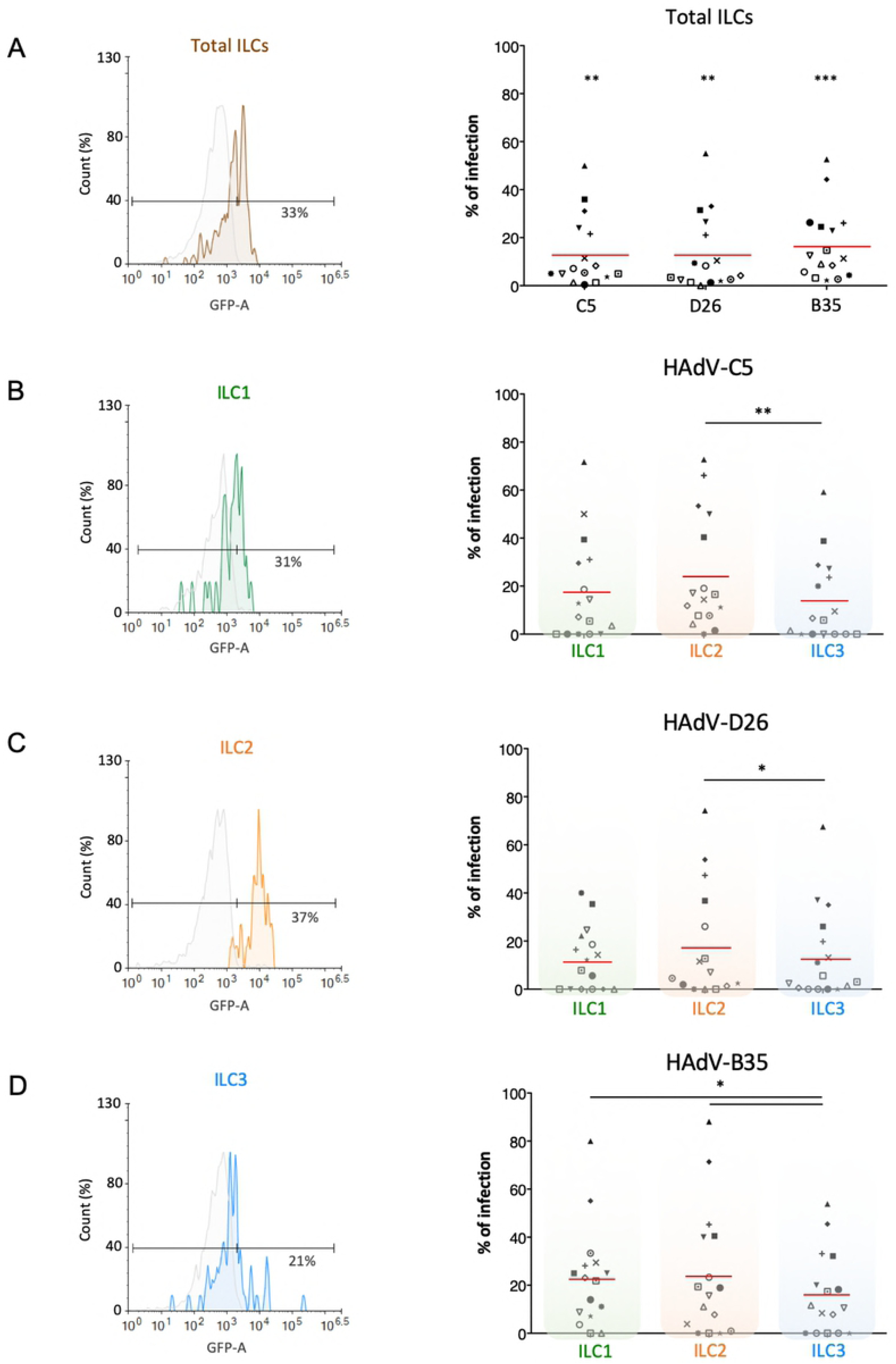
Evaluation of the capacity of ILCs to take up HAdV-C5, -D26 and -B35. HAdV vector-mediated GFP expression in total ILCs and in ILC subsets was quantified 24 h post-incubation (n = 16). The panels on the left (colour-coded to facilitate ILC subset identification) are representative data from a single donor, while panels on the right are cumulative data. **A)** Result from one donor after HAdV-D26 uptake and mean percentages of total ILCs expressing GFP after infection; **B-D)** For each HAdV type, result from one donor after uptake by ILC1, ILC2, or ILC3 and mean percentages of the ILC subsets expressing GFP after infection with HAdV-C5 **(B)**, HAdV-D26 **(C)**, or HAdV-B35 **(D)** (right panel). Statistical analyses were performed using paired Student’s *t* test by comparing uninfected cells and cells challenged with the HAdVs (ns, p > 0.05, * p ≤ 0.05, ** p ≤ 0.01, *** p ≤ 0.001).

### ILCs express receptors used by HAdV-C5, -D26 and -B35

We then screened for the receptors by which HAdVs could be taken up. CAR (coxsackievirus and adenovirus receptor) is a single-pass transmembrane cell adhesion molecule expressed by many cell types and is a primary attachment molecule for numerous HAdV types [35, 36]. We were unable to detect CAR on ILCs (**Fig 3A**). DC-SIGN (or CD209) [37, 38], a C-type lectin receptor present on the surface of macrophages, and conventional and plasmacytoid DCs, is a low affinity/high avidity receptor for some HAdV types [39]. Similar to CAR, we were unable to detect DC-SIGN on ILCs (**Fig 3B**). MHC class I (HLA-ABC) molecules have also been reported to act as a receptor for HAdV-C5 [40], and are high on ILCs (**Fig 3C**). Of note, the diverse haplotypes could be an explanation for the inter-donor variability in HAdV type uptake efficacy. CD46, a type I transmembrane protein that is part of the complement system, is used by some cells to take up HAdV-D26 and -B35 [41, 42]. CD46 was readily detected on essentially all ILCs (**Fig 3D**). We also quantified the level of CD49d (integrin α_4_), which is a low affinity auxiliary receptor for some HAdVs, including HAdV-C5 [43] and was expressed by more than 90% of the ILCs (**Fig 3E**, gMFI for levels can be found in **S3 Fig**). Finally, desmoglein 2 (DSG2), another cell adhesion molecule, is used by some cells to take up some species B and D HAdVs [44]. DSG2 was undetectable on ILCs. Together, these data shed light on the potential mechanism by which ILCs take up HAdV-C5, -D26 and -B35, and are globally consistent with vector-mediated GFP expression.

**Fig 3.**
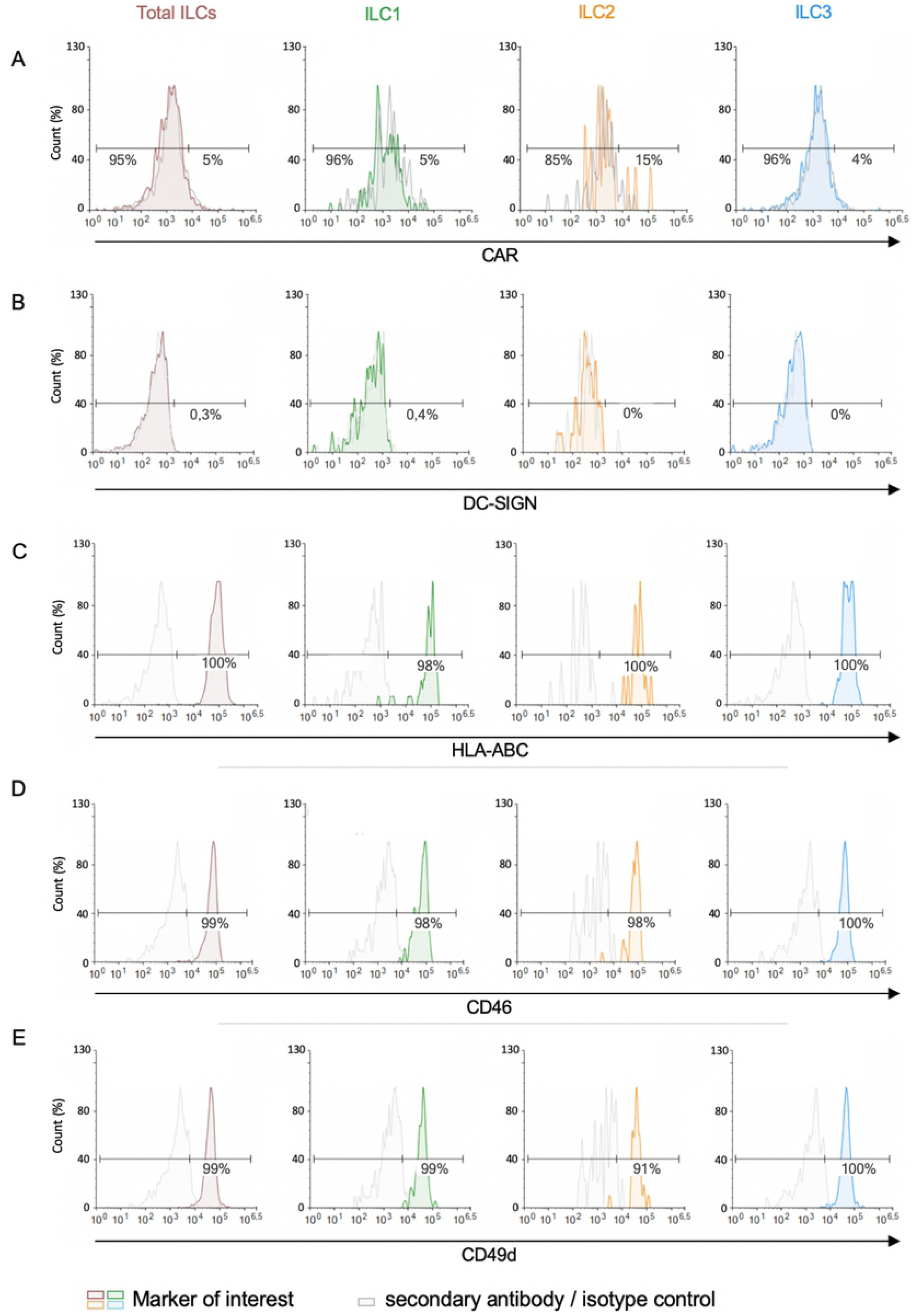
Expression of candidate HAdV receptors by human peripheral blood ILCs. Levels of **A)** CAR, **B)** DC-SIGN, **C)** HLA-ABC (MHC-I), **D)** CD46, and **E)** and CD49d, in freshly isolated total ILCs, ILC1, ILC2 and ILC3. Cell populations were normalised to 100%. Data are representative of 2 - 4 donors.

### ILC phenotypic activation and cytokine secretion after HAdV uptake

ILCs orient adaptive immune responses through the production of cytokines. We therefore examined cytokines involved in antiviral, and initiation or orientation of adaptive immunity following challenge with HAdVs. Due to the limited number of cells/donors, we screened total ILCs. Compared to mock-treated cells, we found an antiviral response consisting of IL- 1β, TNF, IFN-λ_1_, and IFN-γ (<100 pg/ml); IL-8 and INF-β (100 - 200 pg/ml), and IFN-λ_2/3_ (>200 pg/ml), and a Th response consisting of IL-5, IL-6, IL-9, IFN-γ and IL-21 (<100 pg/ml) (**Fig 4A**). TNF and IFN-γ, which have antiviral and Th functions, were comparable in each panel.

**Fig 4.**
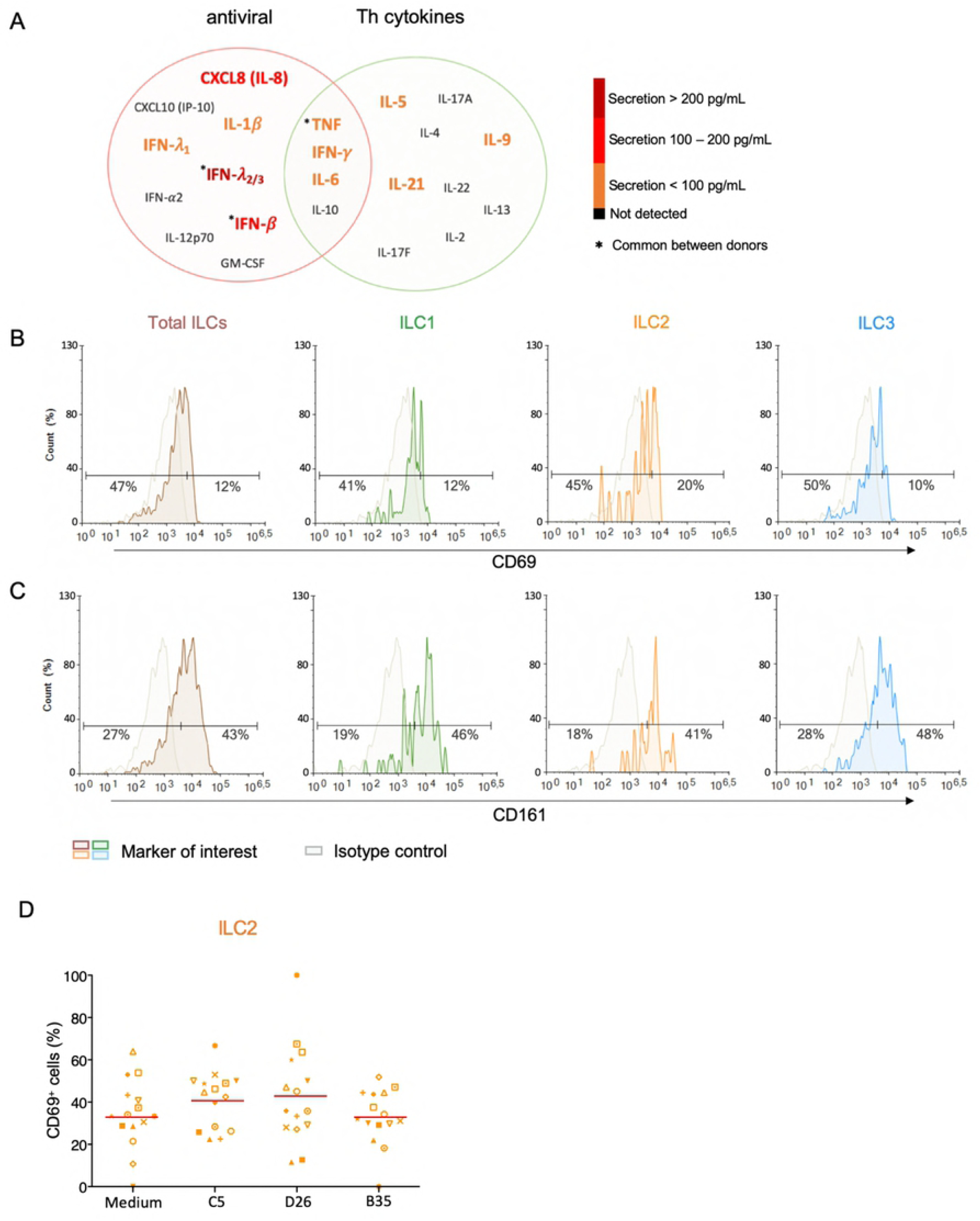
Cytokine release and phenotypic profile of ILCs after HAdV uptake. A) Cytokines belonging to an antiviral panel or a Th screen were quantified from the supernatant of HAdV-challenged ILCs by CBA (n = 5). Cytokine levels are denoted by the colour code and the results were analyzed with the LegendPlexTM software. Only the cytokines whose levels were at least 2-fold higher than that of the controls are shown. Baseline levels of B) CD69 and C) CD161 in freshly isolated total ILCs, ILC1, ILC2 and ILC3 (n > 26). Cell populations were normalised to 100% (n ≥ 5); D) CD69 levels in ILC2 post challenge with HAd-C5, -D26, or B35 (n = 15).

We then characterized the HAdV-induced phenotypic activation. Using ILCs immediately post-enrichment, we quantified the cell surface levels of CD69, an early activation marker expressed by lymphoid cells; CD161, whose expression increases during inflammation (mainly on NK cells); the costimulatory molecules CD80 and CD86; and MHC II molecules. We found that <20% of ILCs had baseline levels of CD69 (**Fig 4B**), while approximately 43% of the cells were CD161^+^ (**Fig 4C**). CD80, CD86, and HLA-DR levels were undetectable (**S4 Fig**). LPS challenge modestly increased the percentage of CD69^+^ ILCs (35%), with the greatest impact on ILC3s (40%). After incubation with HAdV-C5, -D26, or - B35, we observed a selective increase in CD69 levels in ILC2s **(Fig 4D)**. Therefore, ILCs challenged with HAdVs secreted cytokines with antiviral and Th activities and modestly increased a phenotypic marker of activation.

### Impact of pre-existing immunity against HAdVs on the ILC response

HAdV-based vaccines are being used to limit COVID-19 severity and are being trialled for other emerging pathogens [45]. The use of HAdV-C5-based vaccines has shown that, while pre-existing humoral immunity typically reduces vaccine efficacy, vaccine-induced inflammation is not significantly affected [45]. Secondly, a long-term issue will be the ability to reuse HAdV-based vaccines after their nearly global deployment against SARS-CoV-2, which should induce widespread HAdV type-specific immunity. Depending on the cell type and the presence of FcγRs, anti-HAdV antibodies can either inhibit or increase HAdV uptake [46–49]. For example, most sera containing HAdV NAbs are characterized by their ability to inhibit infection of epithelial cells. Yet, these same sera can increase HAdV uptake by professional APCs via FcγRIII (CD16) [34, 50]. FcγR-mediated uptake also increases the phenotypic and functional maturation of moDCs and plasmacytoid DCs [25,33,34,50]. By contrast, Ab that neutralize HAdV-B35 infection of epithelial cells also decreased transgene expression in DCs [51].

Therefore, we asked whether NAbs impact HAdV uptake by ILCs. To form the complexes, we used selected sera that neutralized HAdV-C5, -D26 and -B35 infection of epithelial cells [51]. Following the challenge of ILCs with the HAdV-NAb complexes, we observed a modest increase in cells expressing the transgene when type-specific NAbs were complexed HAdV-C5 and -D26 compared to HAdVs alone (**Fig 5A**). Consistent with previous data, we found that serum that contained HAdV-B35 NAbs tended to decrease the percentage of GFP^+^ ILCs. When broken down into ILC subsets, we observed a modest increase in GFP levels in the presence of NAb-complexed HAdVs for ILC1 and 2 (**Fig 5B & 5C**). However, for ILC3 challenged with HAdV-C5-NAb complexes, we observed a modest decrease in the percentage of GFP^+^ ILCs (**Fig 5D**).

**Fig 5.**
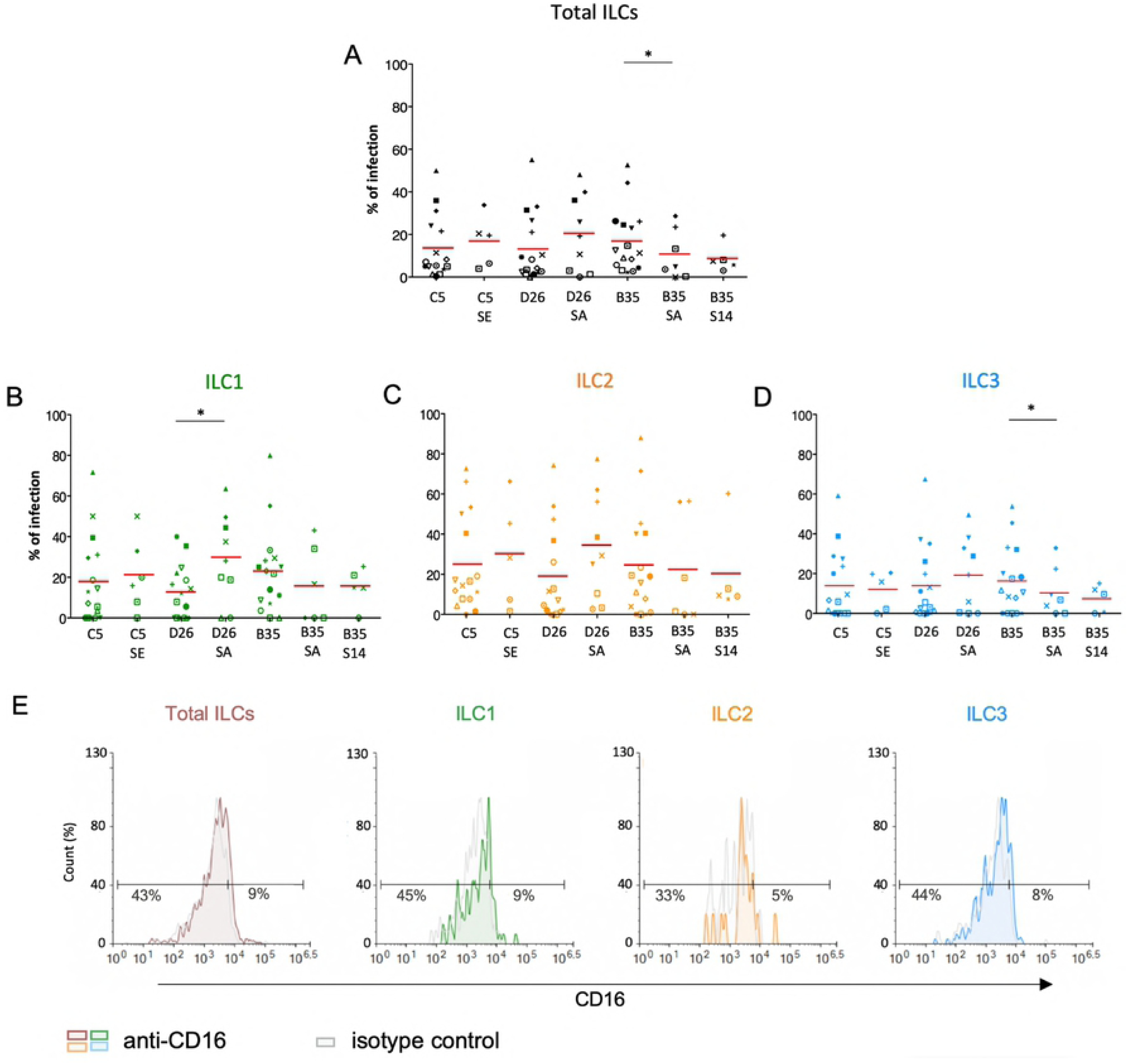
ILC uptake of HAdV-NAb complexes and impact on FcγRIII levels. GFP levels in total ILCs, ILC1, ILC2 and ILC3 24 h post-incubation with each HAdV ± NAbs. Sera “E”, “A”, and “14” inhibit HAdV-C5, -D26, or -B35, respectively, uptake by epithelial cells. **A)** Mean percentages of total ILCs expressing GFP after HAdV ± NAbs challenge. **B-D)** Mean percentages of ILC1, ILC2, and ILC3 expressing GFP after challenge ± NAbs. **E)** Levels of FcγRIII in freshly isolated total ILCs, ILC1, ILC2, and ILC3. Statistical analyses were performed using paired Student’s *t* test by comparing uninfected cells and cells challenged with the HAdVs (ns = p > 0.05, * p ≤ 0.05, ** p ≤ 0.01, *** p ≤ 0.001) (n ≥ 5).

If one compares the subsets, ILC2s (34%) and ILC1s (29%) are more permissive to HAdV- D26-NAb complexes than ILC3s (19%). In addition, ILC1s (15%) appeared to take up more particles/cell (higher gMFI) than ILC3s (7.5%) after HAdV-B35-NAb complex challenge (**S5 Fig**). We therefore quantified the level of FcγRIII, and found lower levels on the surface of ILC1 and 2s versus ILC3s (**Fig 5E**). Globally, ILC2 were the most permissive, while ILC3s appeared the least capable of taking up HAdV complexed with NAbs (summarized in **S5 Fig**).

We then characterized phenotypic activation induced by HAdV-NAbs. In contrast to the HAdVs alone, we found a decrease in CD69 levels following a challenge by NAb-complexed HAdVs (**Fig 6A**). Moreover, each ILC subset tended to have less CD69 on the surface following a challenge by NAb-complexed HAdVs (**Fig 6B**). Together, these data demonstrate that pre-existing B cell immunity can differentially impact the ILC response to HAdVs, likely based on molecules used to take up HAdVs or HAdV-NAb complexes.

**Fig 6.**
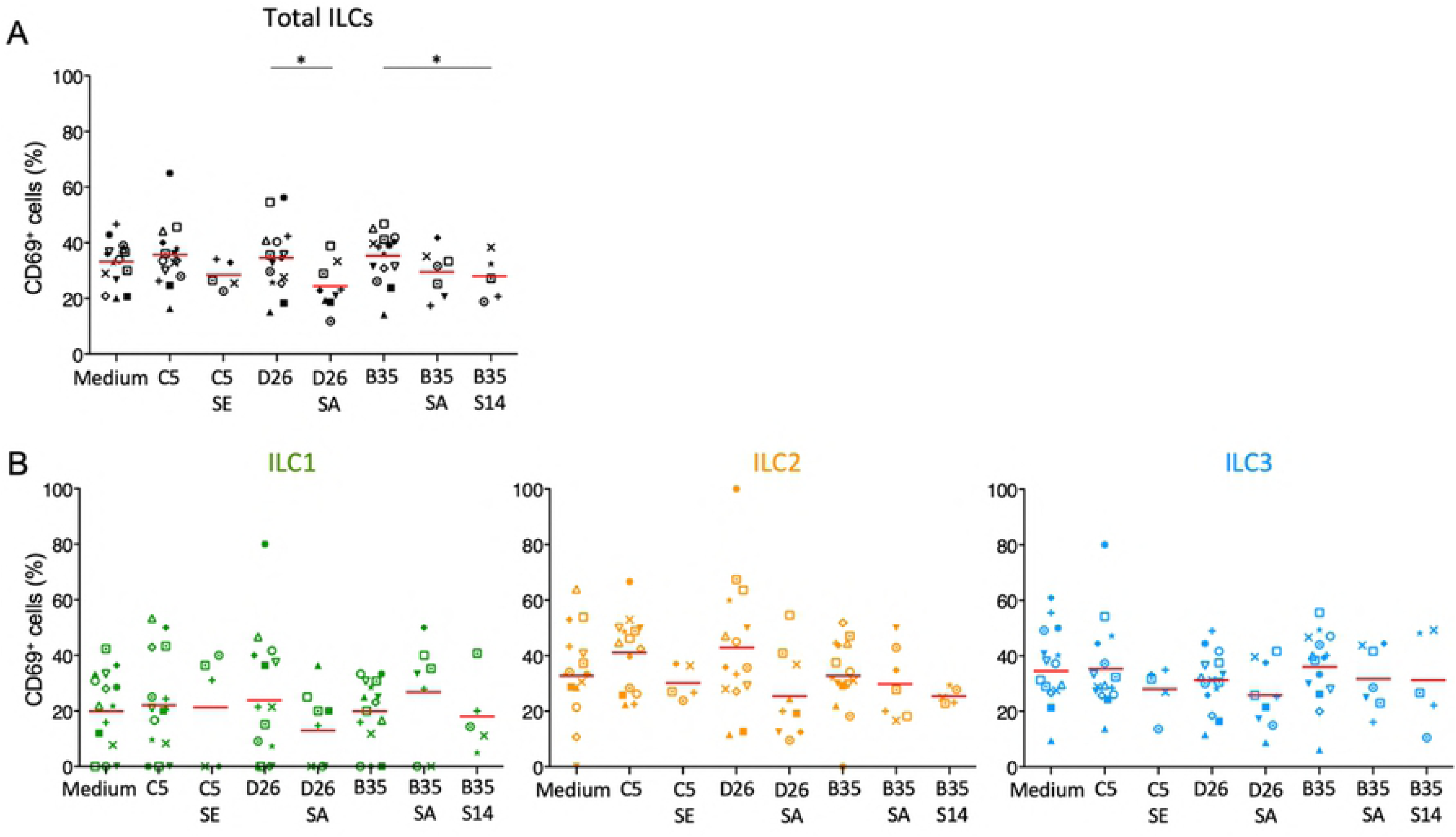
Phenotypic profile by ILCs after NAb-HAdV uptake. CD69 levels were quantified 24 h post-challenge for each HAdV ± NAb complex for **A)** total ILCs and **B)** the ILC subsets (n ≥ 5). Sera “E”, “A”, and “14” inhibit HAdV-C5, -D26, or -B35, respectively, uptake by epithelial cells. Statistical analyses were performed using paired Student’s *t* test by comparing uninfected cells and cells challenged with the HAdVs (ns = p > 0.05, * p ≤ 0.05, ** p ≤ 0.01, *** p ≤ 0.001).

### Pattern recognition receptors

The initial 24 h can be critical when responding to pathogens or vaccines. It was noteworthy that ILC uptake of HAdVs induced IL-8 secretion, which will lead to the recruitment of monocytes and neutrophils (**Fig 4A**). The cytoplasmic content of neutrophils can be as much as 20% HDPs, effector molecules of the innate immune system. Moreover, we previously showed that human neutrophil protein-1 (HNP-1) and lactoferrin bind to HAdV- C5, -D26 and -B35 and, by acting as bridges via TLR4, increase HAdV uptake by DCs [25–28].

As phenotypic and functional activation of innate immunity are initiated by the engagement of PRRs, we screened for the presence of PRRs and markers that serve could as inducers of activation. Despite differences between donors, we found that the majority of the CD3^-^/CD127^+/-^ cells contain relatively low levels of TLR2 (<5%), TLR4 (<7%) and TLR9 (<6%), but significant intracellular levels of TLR3 (**Fig 7A & 7B and S6 Fig**). Importantly though, LPS, a quintessential TLR4 ligand, induces ILCs to secrete pro-inflammatory cytokines, suggesting that while TLR4 levels are not high, TLR4 signalling can be triggered **(S6 Fig**). We therefore asked if HNP-1 or lactoferrin influences ILC uptake of HAdVs. ILCs were incubated with HAdV-HNP-1 or HAdV-lactoferrin complexes and uptake was quantified by GFP expression. In contrast to DCs, we found that the HDP-HAdV complexes either had no effect or were less readily taken up by ILCs (**Fig 7C and S7 Fig**). We then quantified ILC cytokine secretion induced by the HAdV-HDP complexes. When focusing on IL-8 levels, we again found that the response to HAdV-C5 and -D26 separated from that of -B35: when HAdV-C5 and -D26 were incubated with HNP-1 or lactoferrin, ILCs secreted higher levels of IL-8, while HAdV-B35 plus HNP-1 or lactoferrin decreased IL-8 levels compared to the HAdV alone. (**Fig 7D**). The levels of other cytokines did not change notably with respect to the addition of HNP-1 or lactoferrin (**Fig 7E**).

**Fig 7.**
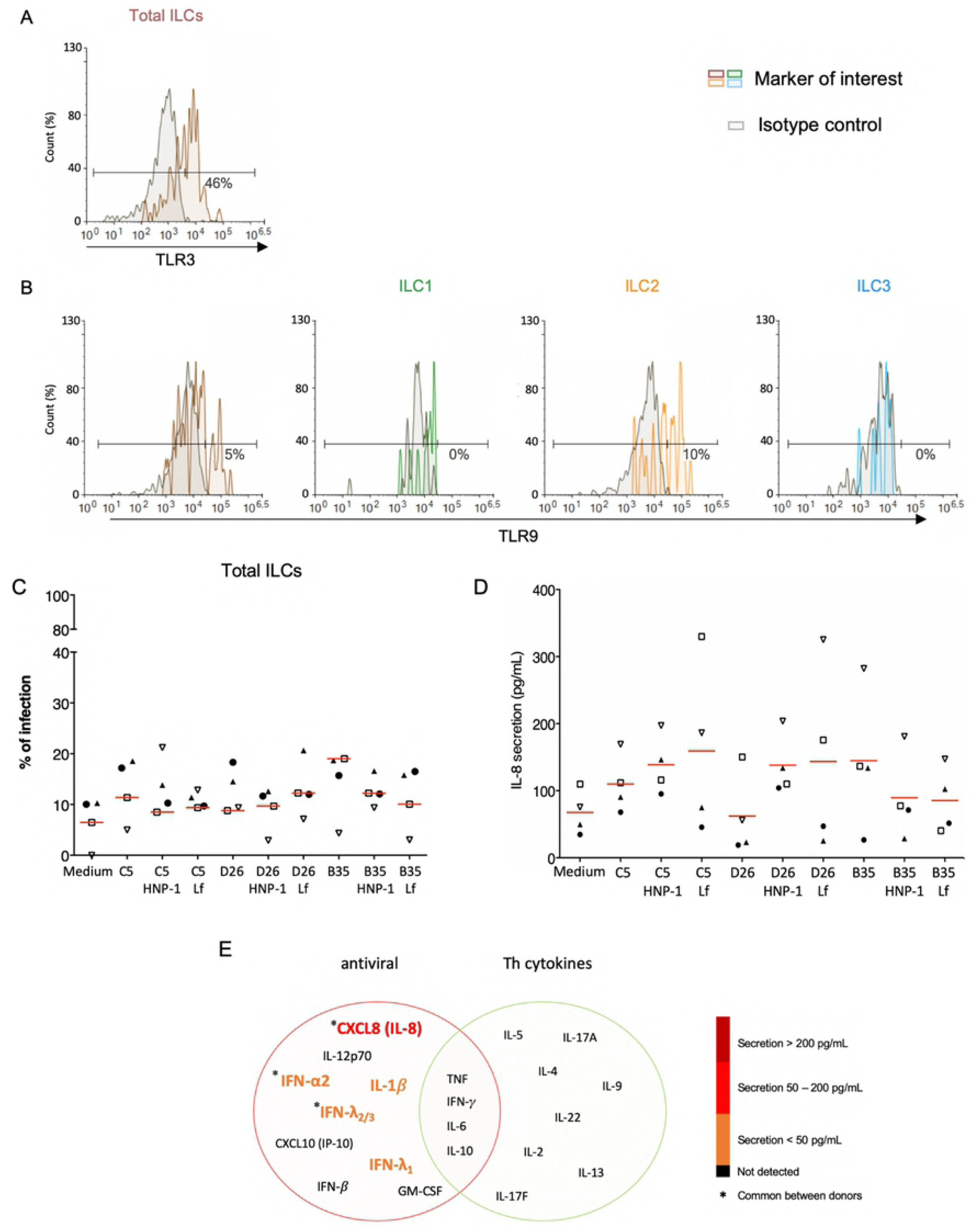
PRR expression and impact of HDPs on uptake and cytokine profile. The levels of **A)** TLR3 by ILCs; and **B)** TLR9 by total ILCs, ILC1, ILC2, and ILC3. Cell populations were normalised to 100%. Results are representative of 2 - 5 donors. **C)** ILC uptake of HAdVs complexed with HNP-1 or lactoferrin: GFP expression by ILCs was quantified 24 h post-challenge with each HAdV ± HDP (HNP-1 or lactoferrin) for total ILCs (n = 4). **D)** Cytokines release in the presence of HNP-1 or lactoferrin: cytokines belonging to an antiviral panel and a Th cytokine panel in the supernatant of HAdV-infected ILCs was quantified using CBA (n = 4). Cytokine levels are denoted by the colour code and the results were analyzed with the LegendPlex^TM^ software. Only the cytokines whose secretion level was at least 2-fold higher than that of the controls are presented. **E)** IL-8 secretion was assessed 24 h post-challenge with each HAdV ± HDP for total ILCs (n = 4). Statistical analyses were performed using paired Student’s *t* test by comparing uninfected cells and cells challenged with the HAdVs (ns = p > 0.05, * p ≤ 0.05, ** p ≤ 0.01, *** p ≤ 0.001).

### Phenotypic maturation and cytokine secretion of bystander ILCs

Because ILCs respond to local cues, we tried to generate an *ex vivo* environment to characterize their response to DCs challenged with HAdVs. In this approach, HAdVs ± NAbs were incubated with moDCs, the moDCs were rinsed to remove HAdVs and NAbs, fresh medium was added, and this latter media was collected 18 h later and added to ILCs. Initially, we observed a 2-3-fold increase in CD69 levels in ILCs due to moDC supernatant **(Fig 8A-C)**. The supernatant from LPS-challenged DCs induced a modest increase in CD69 levels on ILCs, with the greatest impact on ILC3s. When comparing the indirect impact of HAdV-C5, D26 and -B35, it is noteworthy that HAdV-C5, the “gold standard” for HAdV immunogenicity, had the lowest impact on ILC1 and ILC2 phenotypic maturation. When assaying the supernatant from DCs challenged with HAdV-C5-NAb complexes, the number of CD69^+^ ILC1 and 2s increased compared with HAdV-C5 alone. In the case of HAdV-D26- NAb complexes, CD69^+^ levels either decreased (ILC2s, **Fig 8B**) or did not change (ILC1 and 3s, **Fig 8A & 8C)**. Finally, finding serum that neutralizes HAdV-B35 is challenging: in the greater than 400 sera analyzed, we found 1 that inhibited HAdV-B35 infection of epithelial cells. However, due to limited quantities of serum, we were able to perform only 2 assays and therefore the interpretation of these data should take this into account. We found that in contrast to ILC1 and 2s, the ILC3 response to HAdV-B35-NAb complexes was notably higher (**Fig 8C).**

**Fig 8.**
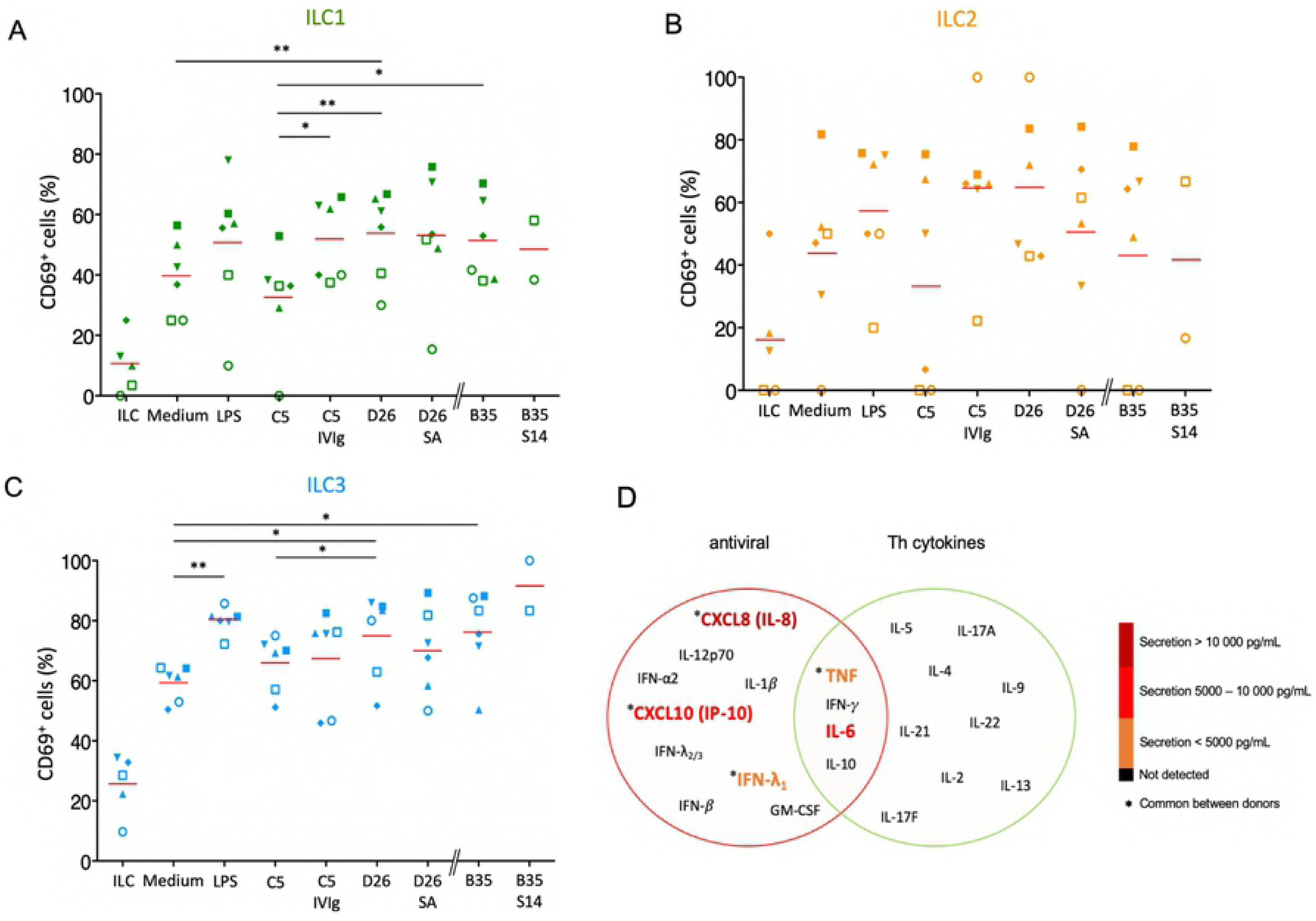
Bystander effect on ILC maturation. We quantified CD69 levels of, and cytokines secreted by, ILCs after indirect challenge moDCs incubated with HAdVs ± NAbs. CD69 levels were measured 24 h post-challenge for each HAdV ± NAbs for total ILCs and different ILC subsets (n ≥ 2). Mean percentages of **A)** total ILCs, **B)** ILC1; **C)**, ILC2, and **D)**, and ILC3, expressing CD69. Statistical analyses were performed using paired Student’s *t* test by comparing uninfected cells and challenged cells (ns p > 0.05, * p ≤ 0.05, ** p ≤ 0.01, *** p ≤ 0.001). **E)** The level of cytokines belonging to an antiviral panel and a Th cytokine panel from bystander ILCs (n = 3). Cytokine levels were denoted by the colour code and results were analyzed with the LegendPlex^TM^ software. Only the cytokines whose secretion level was at least 2-fold higher than that of the controls are shown.

Using the bystander challenge model, we also explored the cytokines secreted by ILCs. As above, we found a mixed antiviral and Th response consisting of TNF, IFN-λ_1_, IL-6, CXCL10, and IL-8 (**Fig 8D and S8 Fig**). We concluded that bystander ILCs respond to DCs challenged with HAdVs ± NAbs and that this response could vary in hosts who have pre- existing HAdV immunity.

## Discussion

The multitasking roles that ILCs play during virus infection have been addressed in numerous situations [10]. However, how pre-existing immunity against a virus, or virus-based vaccines, impacts an ILC response is poorly understood. In this study, we addressed how human ILCs, isolated from peripheral blood, respond to three human adenovirus types, from three species. In addition to the binary ILCs - HAdVs interactions, we investigated how the presence of NAbs and HDPs impacted the ILC response. Because ILC responses are influenced by their environment, we also created an assay to explore how HAdV uptake by antigen-presenting cells (e.g., DCs) influences ILC physiology. We show that *i)* the three ILC subsets can be infected by three HAdV types with variable efficacy; *ii)* depending on the HAdV type, NAbs can either increase or decrease uptake by ILCs; *iii)* ILCs can respond differentially to HAdVs alone or those bound by NAbs or HDPs; and *iv)* the phenotypic profile of, and cytokine release by, ILCs is also responsive to indirect stimulation by HAdV-challenged DCs. Our results demonstrate that the adaptive immune response feeds back into ILC function, which likely impacts HAdV-based vaccine efficacy.

When working with primary ILCs from multiple donors, the heterogeneity of the response is typically considerable. ILC levels in peripheral blood vary with age (up to 7-fold less in older adults compared to children), sex (less abundant in males), and whether responding to a viral infection [11]. While our approach inherently creates challenges for broad interpretations, it nonetheless represents a sampling of the diverse human responses. An important issue to take into account is that we poorly understand how modest changes in the level of cell surface markers of activation, or cytokines secreted impact immune responses. Moreover, using ILCs from peripheral blood creates additional challenges.

Initially, ILCs seed tissues early in the life. During adolescence, it appears that ILCs are replaced by tissue-resident T cells that undertake a role in immune surveillance [52, 53].

Like many respiratory pathogens, the initial HAdV-associated illness is typically a childhood event. HAdVs also cause disease in multiple tissues (eyes, respiratory and gastro-intestinal tracts). The divergent response from tissue resident ILCs should help drive a robust and complex adaptive response to HAdVs. Moreover, in spite of robust anti-HAdV B- and T-cell responses in most adults, HAdVs can maintain long-term persistence [54]. Whether the initial immune response, which likely involves ILCs, plays a role in latency or HAdV-based vaccine efficacy is unknown. Given ILC ability of self-renewal in tissue, it is possible that antigen-specific memory ILCs [55] will, someday, be identified.

Are childhood infections and responses to HAdV-based vaccines in adolescents or adults linked? An argument could be put forward that there is an incompatibility in locales – HAdV-based vaccines are, for the moment, delivered subdermal/intramuscularly, whereas HAdV infections, to the best of our knowledge, rarely occur there. Importantly though, ILCs do not have an obligate tissue-specific residency [56]. ILC homing receptors suggest a context-dependent capacity of some subsets for inter-organ trafficking. In addition, in spite of the T-cell-like functional [8, 9], all ILC subsets took up HAdVs. We also observed variations in the level of potential HAdV receptors, which was consistent with uptake efficacy. These data suggest that ILCs can play a direct role in the initiation of the immune response. Moreover, these data also suggest that functional diversification into Th1, Th2 and Th17/22-like T cells plays a minor role during initial interactions with HAdVs. However, an analysis that we were technically unable to perform was to identify which ILCs secrete which cytokines. ILC uptake of HAdVs generally increased CD69 levels, while the addition of NAbs tended to decrease CD69 level. These observations do not dovetail well with the uptake profile where NAb- HAdV-C5 and -D26 complexes increased the expression of the transgene by ILCs. The challenge of bystander ILCs via infected moDCs also induced a global increase in CD69 levels. The presence of NAbs during moDC challenge increased CD69 levels by ILCs compared to HAdVs alone for -C5 and -B35 and to a decrease for -D26. These patterns underscore the complex role of ILCs after HAdV interactions, particularly in the presence of NAbs.

Yet, as expected, the ILC antiviral response included type I (IFN-β), II (IFN-γ) and III (IFN- λ) IFNs. The levels of TNF, IL-6 and IL-21 were moderate and potentially have a synergistic action rather than individual. TNF is a pro-inflammatory cytokine with a more general role in the induction and stimulation of surrounding immune cells. IL-6 contributes to host defence by stimulating acute phase responses, haematopoiesis and immune responses [57], but has fundamentally different activities depending on its cytokine partners [58]. IL- 21 is a pro-inflammatory, notably inducing IL-8 secretion, maintaining CD8 T cell function and enhancing antigen presentation by phagocytes [59, 60]. In the context of potential HAdV uptake during the first 24 h, the recruitment of HDP-loaded neutrophils could have a significant impact if HNP-1-mediated HAdV uptake influences ILCs directly or indirectly (increasing uptake by local phagocytes) [61]. Unexpectedly, the bystander ILC response induced a cytokine profile similar to that of direct HAdV uptake, but in higher level. We also note the secretion of CXCL10, a pleiotropic molecule that can promote the chemotactic activity of CXCR3^+^ cells, induces apoptosis, and is associated with antiviral responses [62–66].

From these data, we concluded that the ILC response to HAdVs varies in multiple situations **(S9 Fig)**. Moreover, the vast and intriguing inter-donor variability leaves little room for a simple, text-book style conclusion. Identifying biomarkers for ILC status and differences could enable better exploitation and understanding of their responses to viruses and virus-based vaccines.

## Materials & Methods

### Ethics statement

Human blood samples (fresh blood and buffy coat) were obtained from healthy adult anonymous donors at the regional blood bank (EFS, Montpellier, France). The study was approved by the Occitanie & Midi-Pyrenees EFS scientific board (EFS-OCPM: N°21PLER2019-0002). All donors provided written informed consent.

### Adenoviruses

The HAdVs (HAdV-C5, HAdV-D26, HAdV-B35) are ΔE1 making them replication-defective in all cells except E1-transcomplementing cells. HAdV-C5 and HAdV-D26 vectors harbor a GFP expression cassette [67] while HAdV-B35 harbour a GFP variant (YFP) cassette[68]. The vectors were amplified in either human embryonic retinoblasts 911 (HER 911) or 293T E4-pIX cells and then purified to 99% by two density gradients of CsCl [34, 69].

### Enrichment and selection of ILC

Peripheral blood mononuclear cells (PBMC) were isolated on a Ficoll-Histopaque® 1077 gradient (Sigma-Aldrich, Lyon, France). From PMBCs derived from fresh blood, innate lymphoid cells were enriched by negative immunomagnetic selection (EasySep Human Pan-ILC Enrichment Kit, cat# 17975, StemCell). The kit contained a cocktail of magnetic antibodies targeting the major cell lines of the immune system except ILCs. Enrichment was performed according to the manufacturer’s instructions [70]. Freshly enriched ILCs were cultured in a complete medium consisting of RPMI supplemented with 10% human serum AB (sHAB), 10 ng/mL IL-7 (PeproTech®, Neuilly sur Seine, France), 1 mM sodium pyruvate (Gibco) and antibiotics (Penicillin 100 I.U./mL and Streptomycin 100 μg/mL). Different combinations of media were tested in the presence of IL-2, IL-12, and/or IL-7 at different concentrations (at 5, 10, 20, 30, or 90 ng/mL) or a commercial medium specific for NK cells (NK MACS Medium, cat# 130-114-429, MiltenyiBiotec) with lower efficacy. From the ∼60 donors, we obtained, after enrichment, between 2,7 x 10^5^ and 9 x 10^5^ cells/donor, with an average of 6.3 x 10^5^ cells/donor. The percent yield of this enrichment protocol was from 0.02% to 0,7% with a mean of 0,18%. The predicted yield is 0.05 to 0.07% of mononuclear cells.

### Isolation and differentiation of monocytes-derived DCs (moDCs)

From PBMCs derived from buffy coat, monocytes are purified by CD14^+^ expression by positive immunomagnetic selection (MACS system, MiltenyiBiotec). CD14^+^ cells were incubated for 6 days in the presence of 50 ng/ml granulocyte-macrophage colony-stimulating factor (GM-CSF) and 20 ng/ml interleukin-4 (IL-4) (PeproTech®, Neuilly sur Seine, France). The medium used for culture was RPMI, 10% fetal bovine serum (FBS) and penicillin 100 I.U./ml and streptomycin 100 μg/ml.

### Direct infection of ILCs with HAdV vectors

Approximately 2.5 x 10^4^ ILCs in 300 ul of complete medium were incubated in the presence of HAdV-C5-GFP, HAdV-D26-GFP or HAdV-B35-YFP at 10^4^ physical particles (pp)/cell. The medium used for the infection step did not contain human serum. At 6 h post infection, human AB serum (10%) was added to HAdV alone and as NAb complexes. After 24 h of incubation, activation of ILCs was observed via level of the activation markers CD69 and CD161, as well as secretion of antiviral and immunomodulatory cytokines.

### Indirect stimulation of ILCs by moDCs

The first step consists in the stimulation of DCs by HAdVs alone or complexed with NAbs. moDCs (5 x 10^5^ cells in 400 ul of complete medium) were incubated with HAdV-C5-GFP, HAdV-D26-GFP or HAdV-B35-YFP (10^4^ physical particles (pp)/cell) for 6 h in complete RPMI DC medium. 6 h post-infection, the moDC supernatant was discarded and the cells washed with PBS and centrifuged at 1500 rpm for 5 min to remove the HAdVs in the medium. The cells are then cultured in basal ILC medium (RPMI + 10% sHAB + P/S). Approximately 18 h after the medium change (24 h post-infection), the supernatant of activated moDCs was added to freshly isolated ILCs. After 24 h of incubation in the presence of supernatants from stimulated moDCs, ILC activation was characterized by the level of the activation marker CD69, as well as the secretion of antiviral and immunomodulatory cytokines.

### Formation of Ig - HAdV complexes

IVIg® or “Intravenous Immunoglobulin” (Baxter SAS, Guyancourt, France) was used as a control for IC formation and corresponds to a 95% human IgG mix from healthy donor plasma (1,000 from 50,000 donors/batch). IVIg was used in patients with acquired immune deficiencies and autoimmune diseases. For the formation of Ig - HAdV complexes, HAdV- C5-GFP, HAdV-D26-GFP or HAdV-B35-YFP were incubated in the presence of decomplemented sera from a laboratory serum bank for 25 min at room temperature[34]. The sera used may or may not have antibodies specific (or neutralizing antibodies, NAbs) to the different HAdVs used (results of neutralization tests). Serum E (SE) had a very high NAb titer for HAdV-C5 (3500) but no NAbs for HAdV-D26 or -B35. Serum A (SA) had a high titer of NAbs for HAdV-D26 (2500) and low titers for HAdV-C5 (50) and -B35 (0). Serum 14 (S14) had high titers of NAbs for HAdV-B35 (2200) and HAdV-C5 (2000) and a low titer for HAdV-D26 (120). This was the only serum in the range tested (n > 400) for which we detected the presence of a high titer of HAdV-B35 NAbs. After incubation, the newly formed Ig - HAdV complexes were cultured with the different cell types for 24 h. The ILCs (between 2.10^4^ cells and 3.10^4^ cells) or moDCs (5 x 10^5^ cells) were put in the presence of 10^4^ physical particles (pp)/cell of HAdVs.

### Formation of HDP - HAdV complexes

ILCs (between 2 x 10^4^ cells and 3 x 10^4^ cells) were were incubated with 10^4^ physical particles (pp)/cell of HAdVs (in 300 ul of complete medium). For the formation of HDP - HAdV complexes, HAdV-C5-GFP, HAdV-D26-GFP or HAdV-B35-YFP were incubated in the presence of HNP-1 (3.50 ug/mL) or lactoferrin (100 ug/mL) for 30 min at room temperature [26, 27]. These concentrations were chosen to reproduce those found in an inflammatory environment of infected tissues. GFP levels and cytokines release were assessed after 24 h postinfection.

### Flow cytometry

CD127 FITC (cat# 560549, BD Pharmingen) or PE-CF594 (cat# 562397, BD Pharmingen) clone HIL-7R-M21, CD3 PerCP-Cy5. 5 (cat# 560835, BD Pharmingen) or CD3 APC (cat# 300412, BioLegend®) clone UCHT1, CRTH2 (CD294) PE clone BM16 (cat# 563665, BD Pharmingen), CD117 PE-Cy7 clone 104D2 (cat# 339217, BD Pharmingen) were used to identify the population of ILCs and exclude T cells (potential depletion contaminants). CD69 APC (cat# 555533, BD Pharmingen) or FITC (cat# 347823, BD Biosciences) and CD161 PE-Cy5 (cat# 551138, BD Pharmingen) were used to observe activation of ILCs after stimulation. The level of different cellular receptors involved in HAdVs infection was observed by using a panel of antibodies. Anti-CAR (cat# AF336, R&D systems) was used at 1/10th with Donkey anti-goat Alexa Fluor 488 secondary antibody (cat# A11055, Invitrogen), DC-SIGN (CD209) (cat# 561764, BD Pharmingen), Desmoglein 2 FITC (DSG2) clone AH12.2 [71] (cat# sc-80663FITC, Santa Cruz Biotechnology), CD46 APC (targets MCP protein) clone TLA-2-10 (cat# 352405, BioLegend®), CD49d APC (targets α4 integrins) clone 9F10 (cat# 304307, BioLegend®), CD16 (targets FcγRIII) clone 3G8 (cat# 302011, BioLegend®). Several Toll-Like receptors were explored: TLR2 FITC (CD282) clone TL2.1 (cat# 309706, BioLegend®), TLR3 PE (CD283) clone TLR-104 (cat# 315010, BioLegend®), TLR4 APC (CD284) (cat# 130-100-150), MiltenyiBiotec) and TLR9 APC clone eB72-1665 (cat# 560428, BD Pharmigen). The level of HLA-ABC FITC (cat# 557348, BD Pharmigen), HLA-DR FITC (cat# 555811, BD Pharmigen), CD80 FITC (cat# 557226, BD Pharmigen), CD83 FiTC (cat# 556910, BD Pharmingen) and CD86 APC (cat# 555660, BD Pharmingen) was also analysed for ILCs or moDC activation. GFP/YFP expression by infected ILCs or moDCs as well as other previously mentioned markers were assessed by flow cytometry (NovoCyte®). Antibody volumes were adapted to the number of cells for each cell type according to the manufacturer’s instructions. Each cell was incubated for 30 min at 4°C with gentle agitation. The cells were then washed twice (1800 rpm, 4°C, 5 min) and resuspended in 130 uL of buffer (PBS + fetal calf serum). 7-AAD (7-aminoactinomycin D, cat# 559925, BD Pharmingen) was added at 1/250^th^ v/v, 10 min before reading to observe cell viability in each sample. All flow cytometry assays were obtained with the Novocyte® flow cytometer and analyzed with NovoExpress software unless otherwise mentioned.

### Cytometric Beads Assay (CBA)

Supernatants were collected and cytokine secretion was measured using two panels of 13 cytokines (antiviral response and T helper cytokines) by using CBA, a multiplex cytokine quantification (LEGENDplex HU Anti-Virus Response Panel (13-plex) and LEGENDplex HU Th Cytokine Panel (13-plex) (cat# 740390 and cat# 740722, BioLegend) following the manufacturer’s instructions. The concentration of each analyte will be quantified by flow cytometry via the signal intensity and determined using known standard curve and the analysis software provided by the manufacturer (LEGENDplex). We selected the cytokines of interest from a secretion 2-fold higher than control for HAdVs and 2-fold higher than the HAdV alone condition for immune-complexes.

### Statistical analysis

Data were analyzed by GraphPad Prism 5 software. The significance of the results was determined by using Student’s paired *t*-test to make comparisons within each donor.

## Author contributions

Study design & conception: OP, FM and EJK; project direction: FM and EJK. Performed experiments, OP; and analyzed data: OP, FM and EJK. Wrote the manuscript: OP & EJK. Secured funding: EJK.

## Acknowledgements

We thank the imaging facility MRI (ANR-10-INBS-04), Etablissement Français du Sang, and the Plateforme de Vectorologie de Montpellier (PVM, IGMM). We thank Coraline Chéneau for vector preparation, and Anne-Sophie Bedin (UMR 1058, Inserm, Montpellier) for help with the labelling of ILCs. We thank EKL members for constructive comments. We are grateful to Eric Weaver and Andre Lieber for providing HAdV-D26 and HAdV-B35, respectively.

## Supporting information

**S1 Fig. Enrichment, stability and characteristic of human ILCs**

ILCs were identified pre and post negative selection from PBMCs. **A)** Using total PBMCs, we gated on the CD45^+^ population, and within this population, we gated on the Lin^-^ and CD127+ population. **B)** Within the CD127^+^ population, we gated on CRTH2 and CD161 high (ILC2) and low populations (not ILC2). In the not ILC2 population, we screened for CD117 to distinguish between ILC1 and ILC3 (NKp44^-^ and NKp44^+^) populations. ILC1 are identified in fuchsia, ILC2 in light blue and ILC3 in green and dark blue. **C)** Characterization of ILC populations post-negative selection: 85% of the by CD45+ cells were Lin-. Within the Lin- population, >90% were CD127+. **D)** Within the CD127+ population 62%+ were CRTH2 and CD161 high (ILC2). Within the CRTH2 and CD161 low population (not-ILC2s), ∼86 % were ILC3s (NKp44^-^ and NKp44^+^) and 14% were ILC1s. **E)** Quantification of the NK, NKT, T and B cells after negative selection. Lymphoid cells and granulocytes were screened with CD45. In parallel monocytes and macrophages were gated by CD4+ and CD14+. From lymphoid cells, we screened for the presence of CD3 (LyT) and CD19 (LyB), then for CD20 and CD19. Finally, we gated on CD56 and CD16 within lymphoid cells (NKT cells) and within lymphoid cells LyT excluded (NK cells). Summary of NK and NKT cells, LyT and LyB present post-enrichment. Results were obtained with the Navios flow cytometer (Beckman Coulter) and are representative of 3 donors.

**S2 Fig. Infection efficiency of human ILCs with HAdV-C5, -D26 and -B35**

GFP expression by ILCs was measured 24 h post-challenge with each HAdV vector for ILC1, ILC2 and ILC3 (n = 16). The mean percentages of cells expressing GFP after infection were for ILC1: HAdV-C5: 18% / HAdV-D26: 12.5% / HAdV-B35: 23% **(A)**; for ILC2: HAdV- C5: 25% / HAdV-D26: 19% / HAdV-B35: 24% **(B)**; for ILC3: HAdV-C5: 14% / HAdV-D26: 14% / HAdV-B35: 16% **(C)**. Statistics were performed by paired Student’s t test by comparison with uninfected cells and between different HAdV (ns, p > 0.05, * p ≤ 0.05, ** p ≤ 0.01, *** p ≤ 0.001). We noted a significant difference between HAdV-C5 and -D26 and then HAdV-D26 and -B35 for ILC2 (* p ≤ 0.05), but a non-significant difference between the HAdVs tested for ILC1 and ILC3.

**S3 Fig. Median Fluorescence Intensity (MFI) and gMFI of the expression of candidate receptors for HAdVs by human peripheral blood ILCs**

MFI and gMFI for HLA-ABC^+/-^ **(A)**, CD46^+/-^ and CD49d^+/-^ **(B)** cells for each ILC subset and the fold changes were shown for a representative donor. The MFI of 10 (red) correspond to the minimum of fluorescent measured.

**S4 Fig. Activation and co-stimulatory molecule levels in human ILCs**

Levels of CD80 **(A),** CD86 **(B)** and MHC-II (HLA-DR) **(C)** in freshly isolated total ILCs, ILC1, ILC2 and ILC3. Cell populations were normalized to 100%. Results were obtained with the NovoCyte® flow cytometer and analyzed with NovoExpress software. Due to high donor variation, the results presented represent only one donor among the 3 tested (CD80 and CD86).

**S5 Fig. ILC uptake of HAdVs in the presence of NAbs**

**A-C)** GFP expression by ILC1, ILC2 and ILC3 was measured 24 h post-challenge with HAdV-C5-SE **(A)**, -D26-SA **(B)**, or -B35-S14 **(C)** (n ≥ 5). Sera E, A, and 14 have Nabs against HAdV-C5, -D26, and -B35, respectively. **D)** MFI and gMFI for GFP^+^ and GFP^−^ cells for each ILC subset and the fold changes for the mean of 2 representative donors were presented. Statistical analyses by paired Student’s t test were performed by comparison between ILC subsets for each Ig-HAdVs complexes (ns, p > 0.05, * p ≤ 0.05, ** p ≤ 0.01, *** p ≤ 0.001). **E)** Summary of the impact of NAbs on ILC uptake of HAdV-C5, -D26 and - B35.

**S6 Fig. Level of Toll-like receptors by human ILCs directly isolated from peripheral blood**

**A)** TLR2 levels by total ILCs, ILC1, ILC2, and ILC3; **B)** TLR4 levels by total ILCs, ILC1, ILC2, and ILC3. Cell populations were normalized to 100%. Results are representative of 2-5 donors. **C)** Cytokine profile secreted by ILCs after incubation with (LPS) or without (ILCs) LPS for T helper cytokines panels using CBA. Data are representative of one donor.

**S7 Fig. Identification of ILCs and HAdVs uptake in the presence of HDP**

**A-C)** ILCs are among CD3^-^/CD127^+^ lymphoid cells; **D)** GFP levels by ILCs were measured 24 h post-challenge with each HAdV ± HDP (HNP-1 or lactoferrin) for total ILCs.

**S8 Fig. Bystander effect on ILC maturation: IL-8 levels**

IL-8 secretion by human total ILCs was assessed 24 h after indirect challenge by moDCs for each HAdV ± NAbs (n = 3).

**S9 Fig. Summary tables of infection and activation/stimulation of ILCs by HAdVs or infected moDCs in the presence or absence of NAbs**

**A)** Infection capacity of ILCs by HAdVs. **B)** Level of the CD69 marker by ILCs. **C)** Secretion of cytokines by ILCs. **D)** GFP and CD86 levels by DCs after HAdVs challenge. **E)** CD69 level by ILCs after indirect challenge by moDCs. **F)** Secretion of cytokines by ILC after indirect challenge by moDCs. Only cytokines with a secretion level at least 2-fold higher than controls were shown. *IC for Ig – HAdVs complexes*.

## References

1. Spits H, Artis D, Colonna M, Diefenbach A, Di Santo JP, Eberl G, et al. Innate lymphoid cells--a proposal for uniform nomenclature. Nat Rev Immunol. 2013 Feb;13(2):145–9. doi: 10.1038/nri3365

2. Peters CP, Mjösberg JM, Bernink JH, Spits H. Innate lymphoid cells in inflammatory bowel diseases. Immunol Lett. 2016;172:124–31. doi: 10.1016/j.imlet.2015.10.004

3. Kim CH, Hashimoto-Hill S, Kim M. Migration and Tissue Tropism of Innate Lymphoid Cells. Trends Immunol. 2016 Jan;37(1):68–79. doi: 10.1016/j.it.2015.11.003

4. Monticelli LA, Sonnenberg GF, Abt MC, Alenghat T, Ziegler CGK, Doering TA, et al. Innate lymphoid cells promote lung tissue homeostasis following acute influenza virus infection. Nat Immunol. 2011 Nov;12(11):1045–54. doi: 10.1031/ni.2131

5. Britanova L, Diefenbach A. Interplay of innate lymphoid cells and the microbiota. Immunol. Rev. 2017 Sep 1;279(1):36–51. doi: 10.1111/imr.12580

6. Mortha A, Burrows K. Cytokine Networks between Innate Lymphoid Cells and Myeloid Cells. Front Immunol. 2018;9:191. doi: 10.3389/fimmu.2018.00191

7. Golebski K, Ros XR, Nagasawa M, van Tol S, Heesters BA, Aglmous H, et al. IL-1β, IL-23, and TGF-β drive plasticity of human ILC2s towards IL-17-producing ILCs in nasal inflammation. Nat Commun. 2019 14;10(1):2162. doi: 10.1038/s41467-019-09883-7

8. Artis D, Spits H. The biology of innate lymphoid cells. Nature. 2015 Jan 15;517(7534):293–301. doi: 10.1038/nature14189

9. Sciumè G, Shih H-Y, Mikami Y, O’Shea JJ. Epigenomic Views of Innate Lymphoid Cells. Front Immunol. 2017;8:1579. doi: 10.3389/fimmu.2017.01579

10. Vivier E, Artis D, Colonna M, Diefenbach A, Di Santo JP, Eberl G, et al. Innate Lymphoid Cells: 10 Years On. Cell. 2018;174(5):1054–66. doi: 10.1016/j.cell.2018.07.017

11. Silverstein NJ, Wang Y, Manickas-Hill Z, Carbone C, Dauphin A, Boribong BP, et al. Innate lymphoid cells and COVID-19 severity in SARS-CoV-2 infection. Giamarellos-Bourboulis EJ, Rath S, Giamarellos-Bourboulis EJ, Kyriazopoulou E, editors. eLife. 2022 Mar 11;11:e74681. doi: 10.7554/eLife.74681

12. Cooper RJ, Hallett R, Tullo AB, Klapper PE. The epidemiology of adenovirus infections in Greater Manchester, UK 1982-96. Epidemiol Infect. 2000 Oct;125(2):333–45.

13. D’Ambrosio E, Del Grosso N, Chicca A, Midulla M. Neutralizing antibodies against 33 human adenoviruses in normal children in Rome. J Hyg (Lond). 1982 Aug;89(1):155–61.

14. Evans AS. Latent adenovirus infections of the human respiratory tract. Am J Hyg. 1958 May;67(3):256–66. doi: 10.1093/oxfordjournals.aje.a119932

15. Garnett CT, Talekar G, Mahr JA, Huang W, Zhang Y, Ornelles DA, et al. Latent species C adenoviruses in human tonsil tissues. J Virol. 2009 Mar;83(6):2417–28. doi: 10.1128/JVI.02392-08

16. Leung AY-H, Chan M, Cheng VC-C, Yuen K-Y, Kwong Y-L. Quantification of adenovirus in the lower respiratory tract of patients without clinical adenovirus-related respiratory disease. Clin Infect Dis. 2005 May 15;40(10):1541–4. doi: 10.1086/429627

17. Ghebremedhin B. Human adenovirus: Viral pathogen with increasing importance. Eur J Microbiol Immunol (Bp). 2014 Mar;4(1):26–33. doi: 10.1556/EuJMI.4.2014.1.2

18. Mennechet FJD, Paris O, Ouoba AR, Salazar Arenas S, Sirima SB, Takoudjou Dzomo GR, et al. A review of 65 years of human adenovirus seroprevalence. Expert Rev Vaccines. 2019 Jun;18(6):597–613. doi: 10.1080/14760584.2019.1588113

19. Scott MK, Chommanard C, Lu X, Appelgate D, Grenz L, Schneider E, et al. Human Adenovirus Associated with Severe Respiratory Infection, Oregon, USA, 2013–2014. Emerg Infect Dis. 2016 Jun;22(6):1044–51. doi: 10.3201/eid2206.151898

20. Capasso C, Garofalo M, Hirvinen M, Cerullo V. The Evolution of Adenoviral Vectors through Genetic and Chemical Surface Modifications. Viruses. 2014 Feb 17;6(2):832–55. doi: 10.3390/v6020832

21. Zhang C, Zhou D. Adenoviral vector-based strategies against infectious disease and cancer. Hum Vaccin Immunother. 2016 02;12(8):2064–74. doi: 10.1080/21645515.2016.1165908

22. Jönsson F, Kreppel F. Barriers to systemic application of virus-based vectors in gene therapy: lessons from adenovirus type 5. Virus Genes. 2017 Oct;53(5):692–9. doi: 10.1007/s11262-017-1498-z

23. Kumar RK, Foster PS, Rosenberg HF. Respiratory viral infection, epithelial cytokines, and innate lymphoid cells in asthma exacerbations. J Leukoc Biol. 2014 Sep;96(3):391–6. doi: 10.1189/jlb.3RI0314-129R

24. Gregory SM, Nazir SA, Metcalf JP. Implications of the innate immune response to adenovirus and adenoviral vectors. Future Virol. 2011 Mar;6(3):357–74. doi: 10.2217/fvl.11.6

25. Tran TTP, Tran TH, Kremer EJ. IgG-Complexed Adenoviruses Induce Human Plasmacytoid Dendritic Cell Activation and Apoptosis. Viruses. 2021 Aug 27;13(9):1699. doi: 10.3390/v13091699

26. Chéneau C, Eichholz K, Tran TH, Tran TTP, Paris O, Henriquet C, et al. Lactoferrin Retargets Human Adenoviruses to TLR4 to Induce an Abortive NLRP3-Associated Pyroptotic Response in Human Phagocytes. Front Immunol. 2021 May 20;12:685218. doi: 10.3389/fimmu.2021.685218

27. Eichholz K, Tran TH, Chéneau C, Tran TTP, Paris O, Pugniere M, et al. Adenovirus-α-Defensin Complexes Induce NLRP3-Associated Maturation of Human Phagocytes via Toll-Like Receptor 4 Engagement. Banks L, editor. J Virol. 2022 Mar 23;96(6):e01850–21. doi: 10.1128/jvi.01850-21

28. Chéneau C, Kremer EJ. Adenovirus—Extracellular Protein Interactions and Their Impact on Innate Immune Responses by Human Mononuclear Phagocytes. Viruses. 2020 Nov 26;12(12):1351. doi: 10.3390/v12121351

29. Lopez-Lastra S, Masse-Ranson G, Fiquet O, Darche S, Serafini N, Li Y, et al. A functional DC cross talk promotes human ILC homeostasis in humanized mice. Blood Adv. 2017 Apr 11;1(10):601–14. doi: 10.1182/bloodadvances.2017004358

30. Roy S, Jaeson MI, Li Z, Mahboob S, Jackson RJ, Grubor-Bauk B, et al. Viral vector and route of administration determine the ILC and DC profiles responsible for downstream vaccine-specific immune outcomes. Vaccine. 2019 Feb 28;37(10):1266–76. doi: 10.1016/j.vaccine.2019.01.045

31. Parronchi P, Carli MD, Manetti R, Simonelli C, Sampognaro S, Piccinni MP, et al. IL-4 and IFN (alpha and gamma) exert opposite regulatory effects on the development of cytolytic potential by Th1 or Th2 human T cell clones. J Immunol. 1992 Nov 1;149(9):2977–83.

32. Perreau M, Kremer EJ. Frequency, proliferation, and activation of human memory T cells induced by a nonhuman adenovirus. J Virol. 2005 Dec;79(23):14595–605. doi: 10.1128/JVI.79.23.14595-14605.2005

33. Tran TTP, Eichholz K, Amelio P, Moyer C, Nemerow GR, Perreau M, et al. Humoral immune response to adenovirus induce tolerogenic bystander dendritic cells that promote generation of regulatory T cells. Hearing P, editor. PLoS Pathog. 2018 Aug 20;14(8):e1007127. doi: 10.1371/journal.ppat.1007127

34. Eichholz K, Bru T, Tran TTP, Fernandes P, Welles H, Mennechet FJD, et al. Immune-Complexed Adenovirus Induce AIM2-Mediated Pyroptosis in Human Dendritic Cells. PLoS Pathog. 2016;12(9):e1005871. doi: 10.1371/journal.ppat.1005871

35. Loustalot F, Kremer EJ, Salinas S. Membrane Dynamics and Signaling of the Coxsackievirus and Adenovirus Receptor. Int Rev Cell Mol Biol. 2016;322:331–62. doi: 10.1016/bs.ircmb.2015.10.006

36. Bergelson JM, Cunningham JA, Droguett G, Kurt-Jones EA, Krithivas A, Hong JS, et al. Isolation of a Common Receptor for Coxsackie B Viruses and Adenoviruses 2 and 5. Science. 1997 Feb 28;275(5304):1320–3. doi: 10.1126/science.275.5304.1320

37. Maguire CA, Sapinoro R, Girgis N, Rodriguez-Colon SM, Ramirez SH, Williams J, et al. Recombinant Adenovirus Type 5 Vectors That Target DC-SIGN, ChemR23 and αvβ3 Integrin Efficiently Transduce Human Dendritic Cells and Enhance Presentation of Vectored Antigens. Vaccine. 2006 Jan 30;24(5):671–82. doi: 10.1016/j.vaccine.2005.08.038

38. Adams WC, Bond E, Havenga MJE, Holterman L, Goudsmit J, Karlsson Hedestam GB, et al. Adenovirus serotype 5 infects human dendritic cells via a coxsackievirus–adenovirus receptor-independent receptor pathway mediated by lactoferrin and DC-SIGN. J Gen Virol. 2009 Jul;90(Pt 7):1600–10. doi: 10.1099/vir.0.008342-0

39. Korokhov N, de Gruijl TD, Aldrich WA, Triozzi PL, Banerjee PT, Gillies SD, et al. High efficiency transduction of dendritic cells by adenoviral vectors targeted to DC-SIGN. Cancer Biol Ther. 2005 Mar;4(3):289–94. doi: 10.4161/cbt.4.3.1499

40. Hong SS, Karayan L, Tournier J, Curiel DT, Boulanger PA. Adenovirus type 5 fiber knob binds to MHC class I α2 domain at the surface of human epithelial and B lymphoblastoid cells. The EMBO Journal. 1997;16(9):2294–306.

41. Gaggar A, Shayakhmetov DM, Lieber A. CD46 is a cellular receptor for group B adenoviruses. Nat Med. 2003 Nov;9(11):1408–12. doi: 10.1038/nm952

42. Koizumi N, Mizuguchi H, Kondoh M, Fujii M, Hayakawa T, Watanabe Y. Efficient Gene Transfer into Human Trophoblast Cells with Adenovirus Vector Containing Chimeric Type 5 and 35 Fiber Protein. Biol Pharm Bull. 2004;27(12):2046–8. doi: 10.1248/bpb.27.2046

43. Arnberg N. Adenovirus receptors: implications for tropism, treatment and targeting: Adenovirus receptors. Rev Med Virol. 2009 May;19(3):165–78. doi: 10.1002/rmv.612

44. Wang H, Li Z, Yumul R, Lara S, Hemminki A, Fender P, et al. Multimerization of Adenovirus Serotype 3 Fiber Knob Domains Is Required for Efficient Binding of Virus to Desmoglein 2 and Subsequent Opening of Epithelial Junctions. J Virol. 2011 Jul 1;85(13):6390–402. doi: 10.1128/JVI.00514-11

45. Coughlan L, Kremer EJ, Shayakhmetov DM. Adenovirus-based vaccines—a platform for pandemic preparedness against emerging viral pathogens. Mol Ther. 2022 Jan;S152500162200034X. doi: 10.1016/j.ymthe.2022.01.034

46. Bu W, Joyce MG, Nguyen H, Banh DV, Aguilar F, Tariq Z, et al. Immunization with Components of the Viral Fusion Apparatus Elicits Antibodies That Neutralize Epstein-Barr Virus in B Cells and Epithelial Cells. Immunity. 2019 May;50(5):1305–1316.e6. doi: 10.1016/j.immuni.2019.03.010

47. Snijder J, Ortego MS, Weidle C, Stuart AB, Gray MD, McElrath MJ, et al. An Antibody Targeting the Fusion Machinery Neutralizes Dual-Tropic Infection and Defines a Site of Vulnerability on Epstein-Barr Virus. Immunity. 2018 Apr;48(4):799–811.e9. doi: 10.1016/j.immuni.2018.03.026

48. Nimmerjahn F, Ravetch JV. Fcγ receptors as regulators of immune responses. Nat Rev Immunol. 2008 Jan;8(1):34–47. doi: 10.1038/nri2206

49. Guilliams M, Bruhns P, Saeys Y, Hammad H, Lambrecht BN. The function of Fcγ receptors in dendritic cells and macrophages. Nat Rev Immunol. 2014 Feb;14(2):94–108. doi: 10.1038/nri3582

50. Perreau M, Pantaleo G, Kremer EJ. Activation of a dendritic cell-T cell axis by Ad5 immune complexes creates an improved environment for replication of HIV in T cells. J Exp Med. 2008 Nov 24;205(12):2717–25. doi: 10.1084/jem.20081786

51. Ouoba AR, Paris O, Adawaye C, Dzomo GT, Fouda AA, Kania D, et al. Prevalence of Neutralizing Antibodies against Adenoviruses types -C5, -D26 and -B35 used in vaccination platforms, in Healthy and HIV-Infected Adults and Children from Burkina Faso and Chad. 2022 Jun 7;2022.06.07.22276076. doi: 10.1101/2022.06.07.22276076

52. Kotas ME, Locksley RM. Why Innate Lymphoid Cells? Immunity. 2018 Jun;48(6):1081–90. doi: 10.1016/j.immuni.2018.06.002

53. Fan X, Rudensky AY. Hallmarks of Tissue-Resident Lymphocytes. Cell. 2016 Mar;164(6):1198–211. doi: 10.1016/j.cell.2016.02.048

54. King CR, Zhang A, Mymryk JS. The Persistent Mystery of Adenovirus Persistence. Trends in Microbiology. 2016 May;24(5):323–4. doi: 10.1016/j.tim.2016.02.007

55. Martinez-Gonzalez I, Mathä L, Steer CA, Ghaedi M, Poon GFT, Takei F. Allergen-Experienced Group 2 Innate Lymphoid Cells Acquire Memory-like Properties and Enhance Allergic Lung Inflammation. Immunity. 2016 Jul;45(1):198–208. doi: 10.1016/j.immuni.2016.06.017

56. Vivier E, Artis D, Colonna M, Diefenbach A, Di Santo JP, Eberl G, et al. Innate Lymphoid Cells: 10 Years On. Cell. 2018 23;174(5):1054–66. doi: 10.1016/j.cell.2018.07.017

57. Tanaka T, Narazaki M, Masuda K, Kishimoto T. Regulation of IL-6 in Immunity and Diseases. Adv Exp Med Biol. 2016;941:79–88. doi: 10.1007/978-94-024-0921-5_4

58. Kimura A, Kishimoto T. IL-6: regulator of Treg/Th17 balance. Eur J Immunol. 2010 Jul;40(7):1830–5. doi: 10.1002/eji.201040391

59. Elsaesser H, Sauer K, Brooks DG. IL-21 is required to control chronic viral infection. Science. 2009 Jun 19;324(5934):1569–72. doi: 10.1126/science.1174182

60. Parmigiani A, Pallin MF, Schmidtmayerova H, Lichtenheld MG, Pahwa S. Interleukin-21 and cellular activation concurrently induce potent cytotoxic function and promote antiviral activity in human CD8 T cells. Hum Immunol. 2011 Feb;72(2):115–23. doi: 10.1016/j.humimm.2010.10.015

61. Lim AI, Di Santo JP. ILC-poiesis: Ensuring tissue ILC differentiation at the right place and time. Eur J Immunol. 2019;49(1):11–8. doi: 10.1002/eji.201747294

62. Yang J, Richmond A. The angiostatic activity of interferon-inducible protein-10/CXCL10 in human melanoma depends on binding to CXCR3 but not to glycosaminoglycan. Mol Ther. 2004 Jun;9(6):846–55. doi: 10.1016/j.ymthe.2004.01.010

63. Liu M, Guo S, Hibbert JM, Jain V, Singh N, Wilson NO, et al. CXCL10/IP-10 in infectious diseases pathogenesis and potential therapeutic implications. Cytokine Growth Factor Rev. 2011 Jun;22(3):121–30. doi: 10.1016/j.cytogfr.2011.06.001

64. Pandya JM, Lundell A-C, Andersson K, Nordström I, Theander E, Rudin A. Blood chemokine profile in untreated early rheumatoid arthritis: CXCL10 as a disease activity marker. Arthritis Res Ther. 2017 Feb 2;19(1):20. doi: 10.1186/s13075-017-1224-1

65. Spurrell JCL, Wiehler S, Zaheer RS, Sanders SP, Proud D. Human airway epithelial cells produce IP-10 (CXCL10) in vitro and in vivo upon rhinovirus infection. Am J Physiol Lung Cell Mol Physiol. 2005 Jul;289(1):L85–95. doi: 10.1152/ajplung.00397.2004

66. Hayney MS, Henriquez KM, Barnet JH, Ewers T, Champion HM, Flannery S, et al. Serum IFN-γ-induced protein 10 (IP-10) as a biomarker for severity of acute respiratory infection in healthy adults. J Clin Virol. 2017 May;90:32–7. doi: 10.1016/j.jcv.2017.03.003

67. Weaver EA, Barry MA. Low Seroprevalent Species D Adenovirus Vectors as Influenza Vaccines. Miyaji EN, editor. PLoS ONE. 2013 Aug 22;8(8):e73313. doi: 10.1371/journal.pone.0073313

68. Smith JG, Nemerow GR. Mechanism of adenovirus neutralization by Human alpha-defensins. Cell Host Microbe. 2008 Jan 17;3(1):11–9. doi: 10.1016/j.chom.2007.12.001

69. Kremer EJ, Boutin S, Chillon M, Danos O. Canine Adenovirus Vectors: an Alternative for Adenovirus-Mediated Gene Transfer. J Virol. 2000 Jan;74(1):505–12.

70. Valdez Y, Kyei SK, Poon GFT, Kokaji A, Woodside SM, Eaves AC, et al. Efficient Enrichment of Functional ILC Subsets from Human PBMC by Immunomagnetic Selection. The Journal of Immunology. 2018 May 1;200(1 Supplement):51.13–51.13.

71. Kolegraff K, Nava P, Laur O, Parkos CA, Nusrat A. Characterization of full-length and proteolytic cleavage fragments of desmoglein-2 in native human colon and colonic epithelial cell lines. Cell Adh Migr. 2011;5(4):306–14. doi: 10.4161/cam.5.4.16911

